# Persistence of plant-mediated microbial soil legacy effects in soil and inside roots

**DOI:** 10.1101/2020.10.15.340620

**Authors:** S. Emilia Hannula, Robin Heinen, Martine Huberty, Katja Steinauer, Jonathan R. De Long, Renske Jongen, T. Martijn Bezemer

## Abstract

Plant-soil feedbacks are shaped by microbial legacies previous plants leave in the soil. We tested the persistence of such soil legacies after subsequent colonization by the same or other plant species, and whether the microbiome created by the previous plant explains current plant growth. Legacies of previous plants were detectable in soil fungal communities several months after their removal while concomitantly the effect of the current plant amplified in time. Remarkably, bacterial legacies faded away rapidly in the soil and bacterial communities were selected strongly by plant currently growing in the soil. Both fungal and bacterial legacies wrought by the previous plant were conserved inside the root endophytic compartment of the current plant and these endophytes affected significantly the plant growth. Hence, microbial soil legacies present at the time of plant establishment play a vital role in shaping plant growth even as the composition gradually changes in the soil after subsequent plant colonization, as they are taken up as endophytes in the plant. This suggests that plant-soil feedbacks may be partly mediated by a relatively stable endophytic community acquired in early ontogeny while the effects of previous plants detected on soil microbiomes vary between organisms studied. We further show that plants growing in their own soils harbor different endophytic microbiomes than plants growing in soils with legacy of other plants and that especially grasses are sensitive to species specific fungal pathogens while all plant species have less endophytic *Streptomycetes* when growing in their own soil. In conclusion, we show that soil legacies wrought by previous plants can remain present in the soils and inside the roots for months, even when subsequent plants colonize the soil and that these legacies also substantially modulate the plant growth.

## Introduction

Soil microbes are widely acknowledged to be major drivers of plant growth and plant community assembly (1). Plants affect soil microbes via the quantity and quality of rhizodeposits (2,3), and litter (4). These plant-mediated changes in the soil microbiome can influence the growth of other plants that grow later in the same soil (so-called plant-soil feedbacks (5–7)). Plants can negatively influence succeeding plants through accumulation of pathogenic microbes in the soil (8–10), or positively through the build-up of beneficial or mutualistic microbes (11–13). These microbiome-mediated plant-soil feedbacks may be general among functional groups of plants (e.g. grasses and forbs) as these groups markedly differ in their effects on - and sensitivity to - soils. While these microbial soil legacies can have major impacts on plant growth (2), we are still far from understanding and predicting these legacy effects. Specifically, we do not know how persistent soil legacies are, and how the persistence of the microbial legacy effects varies between different plant species that both shape and respond to soil legacies in very different ways is also poorly understood (14,15).

While a ‘current plant’ grows in soil conditioned by a ‘previous plant’, it will respond to soil conditions, but simultaneously also change the microbial legacy in the soil. How these temporal changes contribute to the overall outcome of plant-soil feedbacks is not well understood. Remarkably, however, the specific influence of the current plant on the soil community is a widely held assumption behind plant-soil feedback theories and experiments, whereas the effect of existing soil legacies of the previous plant is often overlooked and lacks rigorous empirical testing. The sensitivity of a plant to the soil microbial community may vary depending on the age of the plant (16–18) and generally, seedlings are considered to be more sensitive than adult plants (19). Freshly germinated seedlings can experience only the soil legacy of the previous plant, whereas older plants on the other hand will experience soils that bear a legacy of previous plants, but which may have also been modified by the current plant. Interestingly, a recent study proposed that the soil microbial community present at the plant germination stage may be a stronger determinant of plant growth than the soil microbial communities that are present at later ontogenetic plant stages (20). Moreover, studies suggest that seedlings are more susceptible to endophytes colonizing the roots than adult plants (21), which may be due to the low levels of chemical defenses in younger plants (19) or their greater need for symbiotic partners to survive.

Endophytic microbes living inside the roots are in closer contact with the plant than the microbes in the soil (22). The endophytes can be beneficial for plant growth through their effects on plant nutrient status, through the protection they provide against pathogens and pests, and via increasing stress tolerance and modulation of plant development (8–10,23,24). Plants inherit endophytic microbes through transfer of microbes from parental plants in seeds (25) but also select their own endophytic microbes from the pool available in the soil (26) and as such, the community structure of endophytes within a plant species is known to differ between soil origins (27,28). It is unknown, however, how the soil legacy created by a previous plant affects the endophytic microbiome of the plant that subsequently colonizes the soil. As most of the endophytes are acquired at early growth stages and often remain in the plant throughout its growth, this suggests that exposure to soil legacies of a previous plant early in life can have long-lasting effects on plant growth, even when these legacies are no longer detectable in the soil surrounding the plant root.

To examine the persistence of plant-specific soil microbial legacies during the next generation of plant growth, we setup a long-term mesocosm common garden experiment with six plant species that are commonly found in former agricultural grasslands and of which plant-soil feedbacks have been well-studied over the past two decades (29,30). As grass species all belong to the same family, but forbs do not, we selected forb species from one family as well. The three selected grasses were *Alopecurus pratensis, Festuca ovina* and *Holcus lanatus* (all Poaceae) and the three selected forbs were *Hypochaeris radicata, Jacobaea vulgaris* and *Taraxacum officinale* (all Asteraceae). We first created six distinct soil microbial legacies by growing the plants as monocultures in 200-L mesocosms for 12 months (31). We then divided each mesocosm into six physically separated sections, in which we planted all the six responding plant species (see Figure 1 for set-up). We monitored the soil microbiome in each section by non-destructive repeated sampling for five months and examined changes in the microbiome (bacteria and fungi) caused by the previous and the current plant over time. After five months of plant growth, we destructively harvested the plants to examine their responses to the soil legacies and analysed the root endophytic microbiome (22,28).

**Figure 1.**
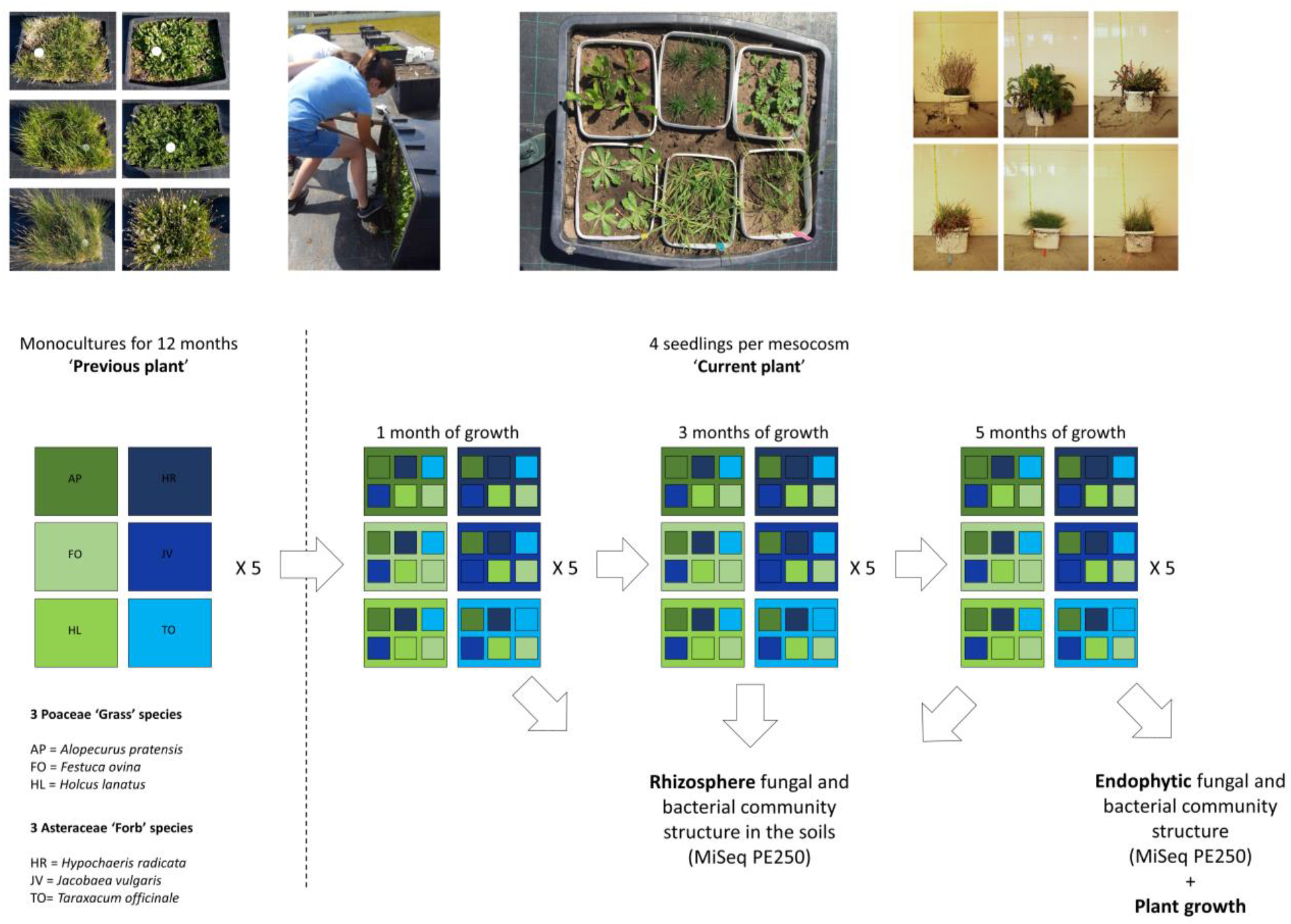
Set-up of the experiment. In short, monocultures of six plant species were maintained for 12 months in (6 x 5) 30 mesososms. After that all mesocosms were flipped to their side and top soil was divided into six smaller mesocosms. See supplementary video for how this was done. The existing plants were removed in mid-May and after three weeks (in June) four seedlings of one of the same six plants were planted in each small container reciprocally. This equaled to (30 mesocosms x 6 plants) 120 smaller mesocosms. Soil from each of these mesocosms were sampled one month (July), three months (September) and five months (November) after planting the plants. Soil was sampled so that four soil cores were taken per mesocosm next to each plant and combined into one composite sample per small mesocosm. After five months, the containers were destructively harvested, plant biomass measured from dried and washed plant material, and endophytic microbial communities were surveyed from sterilized root samples.

We tested the following hypotheses: (i) Plants create directional changes in soil microbiomes and these differ between the grasses and forbs, and between plant species (Figure 2A). (ii) The soil microbial legacy of the previous plant diminishes over time, while the effects of the current plant on the soil microbial community concomitantly increase with time (Figure 2B). We further hypothesize (iii) if the endophytes are acquired by plants in early growth stages, the effect of the previous plant, even if not detected anymore in the soil, will still be visible in the endophytic root microbiomes. We also test if plant growth is related to soil microbial community composition and expect (iv) that endophytic microbes have stronger relationships with plant growth than soil and rhizosphere microbes (22,24,32). Lastly, (v) we tested whether conspecific negative feedbacks (i.e. poor performance in soil from their own species, compared to other soils (11,15)), are due to an accumulation of species-specific pathogens in the rhizosphere (33), and we investigate at which level this operates. To achieve these aims, we analysed both bacterial and fungal communities in the soils at three time points and inside the roots, and relate the community composition of microbes in soils conditioned with different plant species to the plant biomass of the next plant at the end of the experiment. We expect based on previous work that bacterial soil legacies will have a faster turnover time than fungal legacies (31,34) due to differences in growth strategies/traits (35), and hence that the legacy effects of the preceding plant will be detected for longer time periods in the fungal than in the bacterial community.

**Figure 2.**
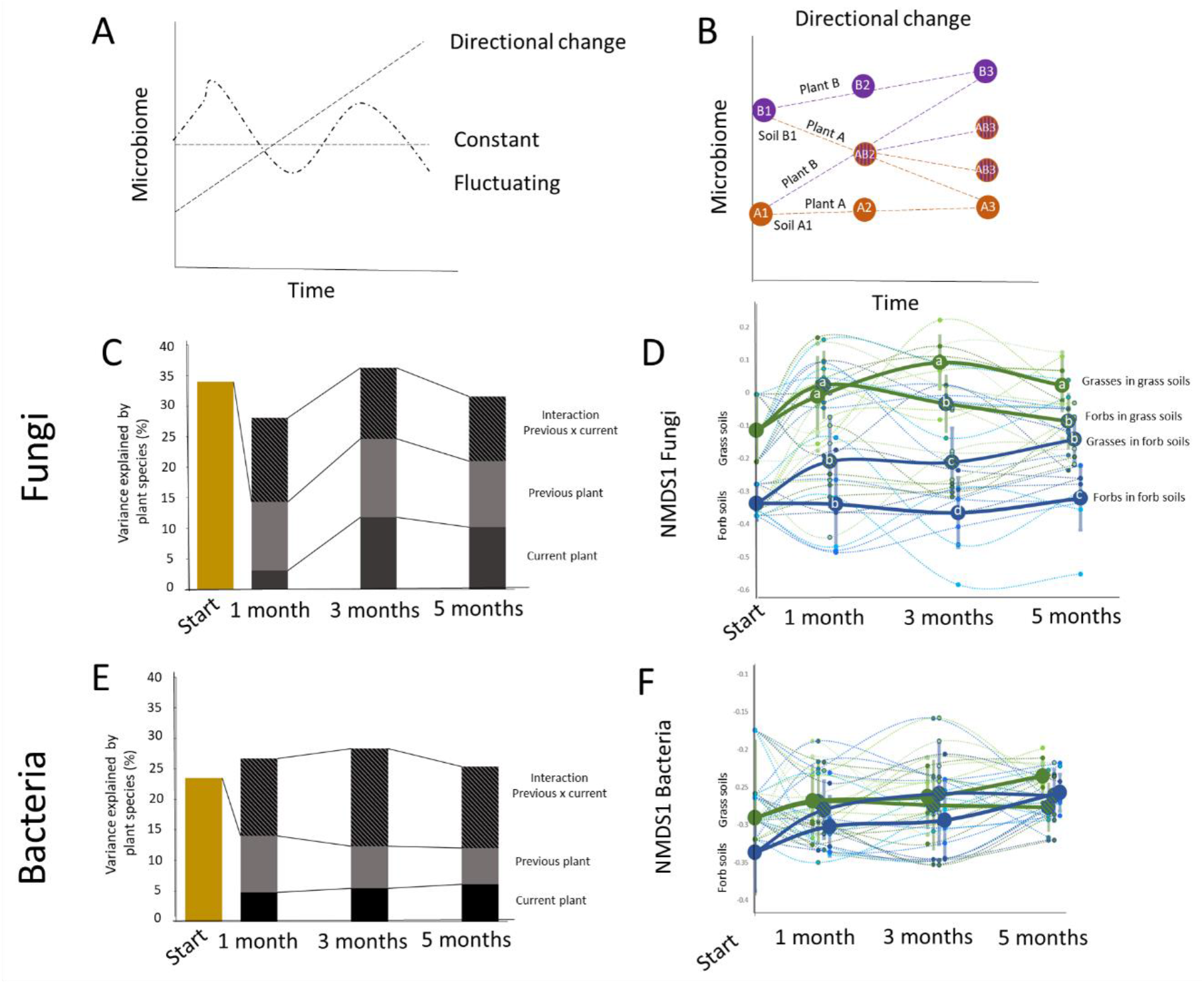
Theoretical and observed change in microbiomes in time. Theoretical framework and observed effects of conditioning (previous) and responding (current) plant on soil microbial community composition. (A) we expected that the microbiome would change in time under a new plant community in either directional, constant, or fluctuating way. (B) Furthermore, we expected that the changes would lead to convergence in microbiomes and that in time the responding microbiome would create its own microbiome type and that this would be depending on the initial microbiome and hence the plant in the conditioning phase. Variance in fungal (C) and bacterial (E) community structure in soils explained by current and previous plant species and their interaction in the beginning (explained by the current plant species) of the experiment and after 1, 3, and 5 months. Changes in community structure of fungi (D; depicted as NMDS1, for full data see supplementary figure 3) and of bacteria (F; depicted as NMDS1, for full data see supplementary figure 4) in time and per plant species (thin lines) and averaged per plant functional group (thick lines). The colors refer to plant species, green colors mark grasses and blue colors forbs. Error bars of points on the thick line depict the standard error between plant species. The letters in the circles note statistical significance.

## Results

### Directional changes in soil microbiomes

First we investigated the direction of temporal changes in microbiomes due to current and previous plant species and plant families (Fig. 2A & B). After one year of plant growth, each plant species had created its unique microbiome (PerMANOVA R^2^=0.35 for fungi and R^2^=0.24 for bacteria, for both p<0.001; Fig. 2 C & E) and the soil microbiomes also differed significantly between the two plant families supporting hypothesis i) (PerMANOVA R^2^=0.13 for fungi and R^2^=0.07 for bacteria, for both p<0.001; Fig. 2 D & F, Fig. S1 & S2). The effects of the previous and current plant family on bacterial and fungal communities in the soil changed over time and the pattern followed hypothesis ii), but only partly. One month after planting the current species, for both the fungi and bacteria, a larger part of the variation in the soil microbial community structure was explained by the identity of the previous plant than by that of the current plant (for bacteria: PerMANOVA *previous* R^2^=0.09 and *current* R^2^=0.05, for both p<0.001 and for fungi PerMANOVA *previous* R^2^=0.11, p<0.001 and *current* R^2^=0.03, p=0.11; Fig. 2 C & E, Fig S1-S3). For soil bacteria, the effects of the previous plant diminished over time, while the effects of the current plant increased (Fig. 2E). For soil fungi, however, the amount of variation in community composition that was explained by the legacy of the previous plant species was highest three months after planting the new species (PerMANOVA R^2^=0.13, p<0.001), and five months after planting, this effect was still present and larger than the amount of variation explained by the current plant species (PerMANOVA R^2^=0.11, p<0.001; Fig. 2C).

Interestingly, especially for fungi, the family of the previous and current plant had a large effect on community assembly. Irrespective of the plant species identity, the Poacaeae (grasses) and Asteraceae (forbs) left distinct fungal legacies and also responded differently to these legacies (Fig. 2 D; Fig. S3). After one month most variation in fungal communities in the soil was explained by whether the plant was growing in soil with a legacy of a Poacaeae or of an Asteraceae (PerMANOVA R^2^=0.08, p<0.001; ANOVA on NMDS1: F_1,34_=41.75, p<0.001, Fig. 2D, Fig S1, Fig. S3). Two months later, we detected a significant effect of the current plant family (PerMANOVA R^2^=0.05; ANOVA on NMDS1: F_1,34_=97.60, for both p<0.001), but also of the family the previous plant belonged to (PerMANOVA R^2^=0.08; ANOVA on NMDS1: F_1,34_=18.71, for both p<0.001). Each current/previous family combination created a unique mycobiome type (ANOVA on NMDS1: F_3,32_=38.84, p<0.001; Fig 2D) so that soils with forbs currently growing in them were moving closer to soils with a legacy of forbs, while soils with grasses currently growing in them more closely moved towards soils with a grass legacy. After five months of growth, we could distinguish three types of mycobiomes, as soils with a forb legacy and with currently grasses and soils with a grass legacy with current forbs had diverged and became similar (ANOVA on NMDS1: F_3,32_=26.42, p<0.001; Fig 2D).

For bacteria, the soil microbiomes of the two families differed less, and for the first two time points, no effect of previous or current family was detected (Fig. 2F). However, after five months of conditioning, bacterial communities differed between soils with current grasses and forbs, but only in the soils with a legacy of grasses (ANOVA on NMDS1: F_3,32_=2.97, p<0.05; Fig 2D). Importantly, the time point of sampling, affected the bacterial community composition much more strongly (PerMANOVA R^2^=0.39, p<0.001; Fig. S2) than the fungal composition (PerMANOVA R^2^=0.09, p<0.001; Figure S1) indicating that the bacterial community is less temporally stable than fungal community (Fig. 2A)

### Effect of soil and its microbiome on plant growth

We destructively harvested the current plants at five months, to investigate if plant growth was affected by previous plant-mediated soil legacies. Aboveground biomass of all plant species was affected by the soil legacies wrought by the previous plant (Fig 3A-B). For grasses, aboveground biomass in soils with a grass legacy was lower than in soils with a forb legacy (ANOVA F_1,24_=19.52, p<0.001; Fig 3A), and for the three forb species, a negative effect of growing in their own soil was detected (Fig. 3A). Across all plant species, aboveground biomass at harvest was related to the soil fungal community structure at all the time points measured (Fig. 3C), while no relationship was observed between the bacterial community structure in the soil and aboveground plant biomass at harvest (Fig. 3D). The effect of the soil fungal community on plant shoot biomass at harvest was detected at all the time points and the effect was strongest after five months of plant growth (ENVFIT: R^2^=0.06, p<0.001; Fig. 3E) which was related also to the first axis of a multivariate NMDS ordination for the fungal community (Pearson: R^2^=0.07, p<0.001). When the effects of the fungal community at the time of harvest (i.e. after five months) on plant growth were evaluated for each test plant species separately, we detected that the aboveground biomass of all grass species was explained by soil fungal community composition, but for the forbs, only the shoot biomass of *Taraxacum officinale* (TO) was weakly related to soil fungal community composition (Fig. 3F).

**Figure 3.**
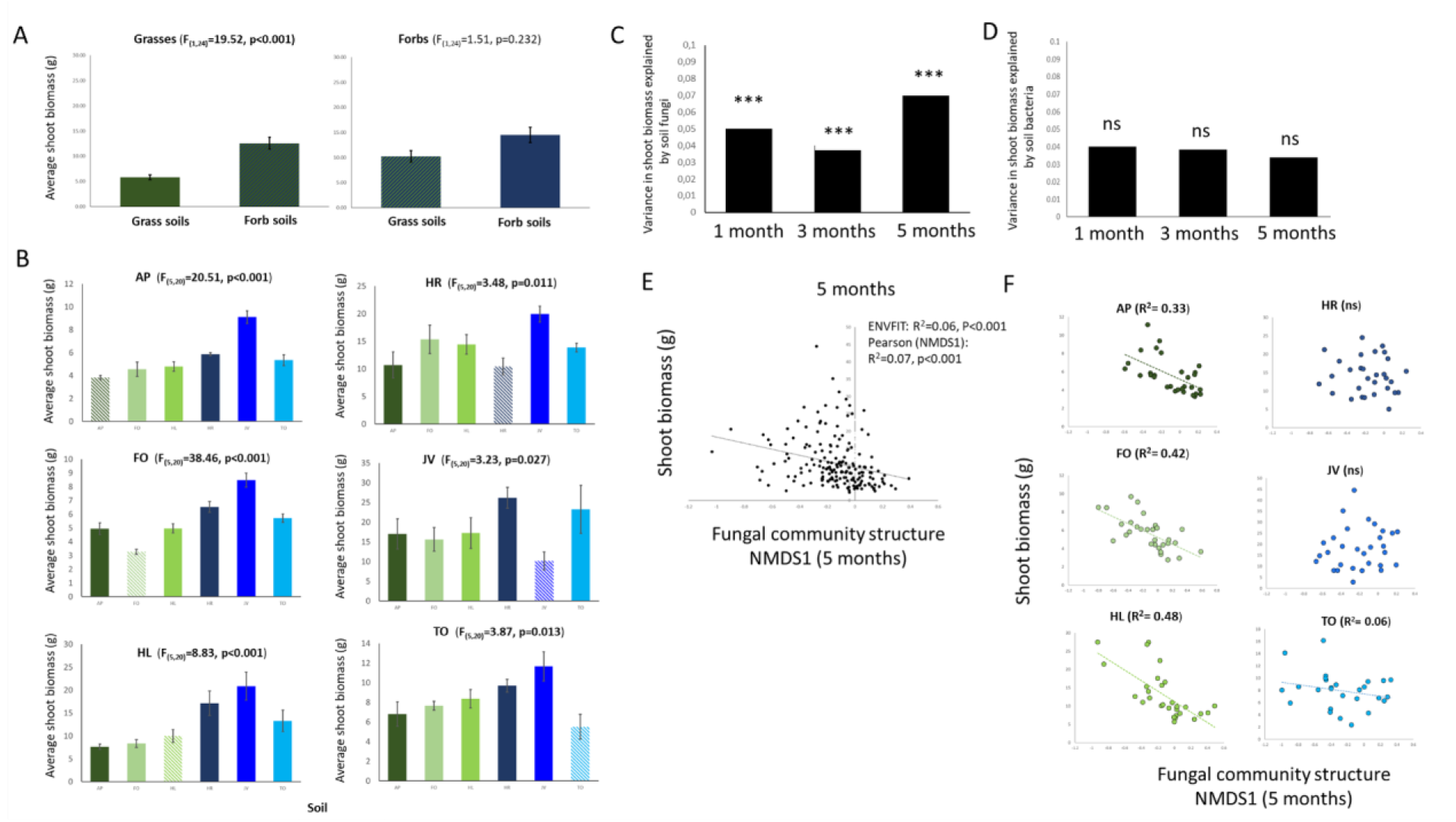
Plant growth across soils and soil microbial communities explaining the plant growth. Plant growth in the feedback phase affected by previous plant and explained by soil microbial communities. Average shoot biomass (dw) of the responding grasses and forbs grown in grass and forb soils (A) and of individual plants grown in all the possible soils (B). Own soil functional group and species (conspecific feedbacks) are shown with striped bars while other soils in full colors. Grasses are depicted with green colors and forbs with blue. Statistical values are shown above each graph for the full model. Bars depict mean values with standard error. The variance in plant shoot biomass (dw) explained by soil fungal (C) and bacterial (D) community composition at different time points. The statistical results given in panels C and E are PERMANOVAs on Bray-Curtis distances and based on full data. (D) Relationship between shoot biomass and fungal community structure measured by NMDS1 (Pearson correlation) and by using all the data and full model (ENVFIT), and (F) divided per current plant species. In (F) correlation coefficients are shown only for plant species significantly responding to changes in fungal community.

### Legacy effects on endophytic microbes

To evaluate the hypothesis (iii) that the previous plant influences the endophytic root microbiome of the current plant, we examined the effect of the previous and current plant species on the fungal and bacterial communities inside plant roots. The two families to which the previous plant belonged significantly differed in how they influenced endophytic fungi (Fig. 4A) and endophytic bacteria (Fig. 4B) in the roots of the current plant. Furthermore, there was an interaction between the family of the previous plant and the family of the current plant on endophytic community structures (Fig. 4 A & B). For both endophytic bacteria and fungi, the identity of the current plant explained most of the variation in community structure (PerMANOVA R^2^=0.48 for bacteria and R^2^=0.26 for fungi, for both p<0.001; Fig. 4C). However, the legacy of the previous plant also significantly affected the composition of the bacterial and fungal root endophytic communities (PerMANOVA R^2^=0.06 for bacteria and R^2^=0.09 for fungi, for both p<0.001; Fig. 4C), and there was a significant interaction between the legacy of the previous plant and the identity of the current plant (R^2^=0.09 and R^2^=0.15, for bacteria and fungi, for both p<0.005; Fig. 4C). For fungi, the effect size of the previous plant family was similar in the soils and in the roots after five months of growth (R^2^=0.09 in the roots and R^2^=0.06 in the soil) while for bacteria the effect of the family of the previous plant was stronger in the roots than in the soil (R^2^=0.07 in the roots and R^2^=0.02 in the soil).

Different endophytic fungal and bacterial groups were selected for by current plant species that were growing in soil with a legacy of previous plant species (Fig. 4D). Bacterial phyla like *Actinobacteria, Patescibacteria*, and *Fibrobacteres* were strongly affected by current plant species (LME: F values > 20, p<0.001 after FDR correction), while especially fungal classes such as Magnaporthales and Sebacinales and also bacterial phyla such as *Acidobacteria* and *Nitrospirae* were affected by both the previous and the current plant (LME: F values for previous plant >5, p<0.001 after FDR correction; Fig. 4D).

**Figure 4.**
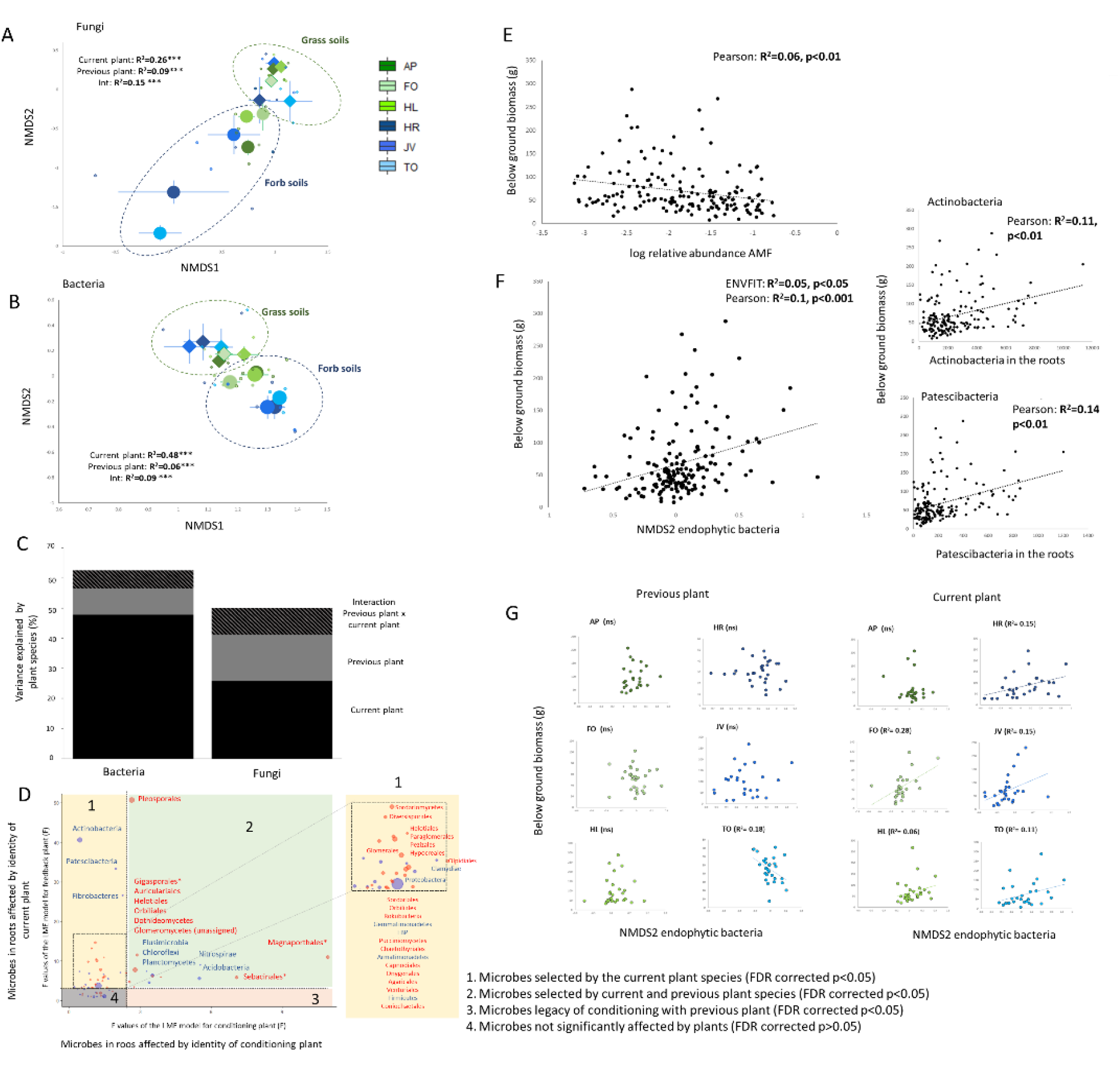
Endophytic microbes and their effects on plant growth. Fungal and bacterial root endophytes affected by conditioning and responding plant species and functional groups and their relationship with responding plant growth. (A & B) NMDS using Bray-Curtis distance on the effect of current plant (colours; green colours grasses, blue colours forbs) and previous plant functional group (shapes; circles represent forb soils and triangles grass soils) on fungal (A) and bacterial (B) community structure. (C) variance in root endophytic bacterial and fungal community structures explained by conditioning (previous) and responding (current) plant and their interaction. (D) endophytic fungal orders (red) and bacterial phyla (blue) whose abundances are significantly (FDR p<0.05) changed by conditioning plant species (3, orange), responding plant species (1, yellow) and by both (2, green). The dashed lines indicate the minimum F values that are significantly explaining the LME model after FDR correction. (E) the relationship between (log relative abundance of) AMF inside the roots and plant below ground biomass. (F) relationship between root biomass and bacterial community structure measured by NMDS2 (Pearson correlation) and by using all the data and full model (ENVFIT), and read numbers of two bacterial phyla (Actinobacteria and Patescibacteria) most strongly correlated with belowground biomass. (G) Relationship between endophytic bacterial community structure measured with NMDS2 and belowground plant biomass divided per conditioning soil and responding plant. Only for significant correlations, the correlation coefficient is shown.

In order to answer hypothesis iv) we related the endophytic community composition further to the plant growth parameters. The relative abundance of arbuscular mycorrhizal fungi inside the roots was associated with root biomass across plant species but this relationship was negative (Pearson: R^2^=0.06, p<0.01; Fig. 4E). Root biomass was also significantly related to bacterial community composition inside the roots (ENVFIT: R^2^=0.05, p<0.05; Fig. 4F). Specifically, the relative abundance of *Actinobacteria* and *Patescibacteria* inside the roots correlated with greater root biomass (Fig. 4F). For all but one of the current plant species, there was a significant relationship between endophytic bacterial community and root biomass, but the magnitude of the effect (measured as R^2^) varied among species (Fig. 4G).

### Microbiome effects on plant growth on own soils

Lastly, we tested hypothesis (v) on the role of microbiome and individual microbes in regulation of conspecific feedback, i.e., plants growing on soil legacies created by their own species. On average, all plant species used in this experiment performed worse in soils with a legacy of the same plant species than in soils from other plant species (Fig. 3B, Fig. 5A). Three plant species (*Holcus lanatus, Jacobaea vulgaris*, and *Taraxacum officinale*) had a significantly different fungal composition when grown in the soils where same plant species was grown earlier than in soils with other plant species one month after planting (PERMANOVA: R^2^>0.05, p<0.01; Fig. 5B & S4). For *Holcus lanatus*, this effect was related to the larger relative abundance of fungal potential plant pathogens in its own soil than in other soils in the first time point (F=8.63, p=0.006: Fig. S6), while for *Jacobaea vulgaris* we detected a reduction of AMF in its own soil compared to other soils at the first time point (F=6.65, p=0.016: Fig. S6). For the other species (*Alopecurus pratensis*, *Festuca ovina*, and *Hypochaeris radicata*), a similar effect was observed for the soil samples collected at five months (Fig. 5B, Fig. S4). We also detected differences in bacterial communities when growing in their own soil, especially for *J. vulgaris*, *F. ovina* and *H. radicata*, after three and five months of growth (Fig. 5B, Fig. S5). For the endophytic fungi, we detected significant effects of growing in own soil soil for *A. pratense, F. ovina* and *J. vulgaris*, while endophytic bacteria differed between own soil and other soils for *A. pratense, F. ovina* and *H. radicata* (Fig 5B, ‘roots’).

**Figure 5.**
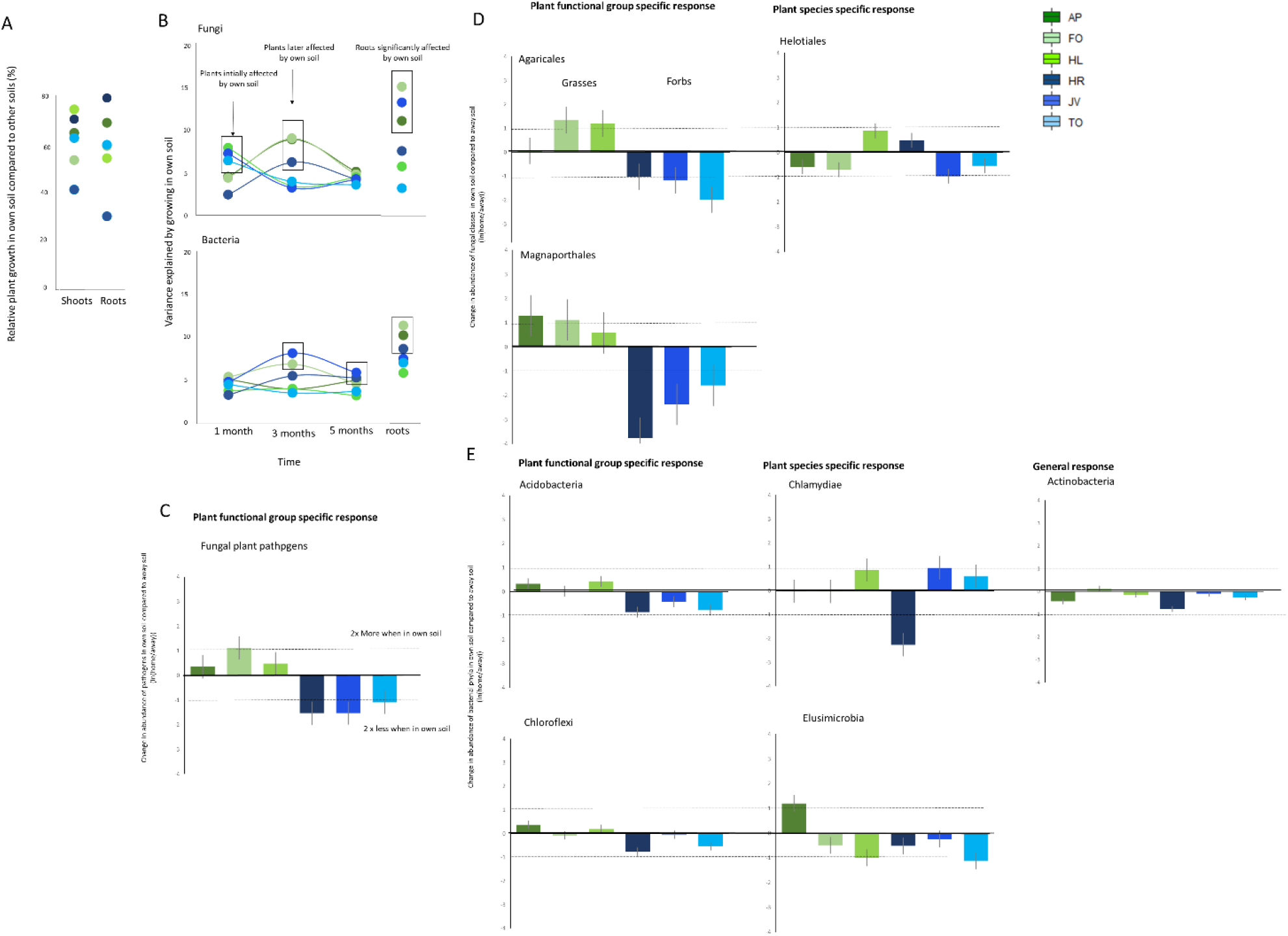
Home-away effects on plants and microbial groups. The effects for plants to grow on their own vs other soils. (A) the relative plant growth of the plant in its own soil compared to other soils for shoots and roots. (B) the variation in variance explained by growing in own soil (vs other soils) in time and between plants species estimated with PERMANOVA. Significant effects are circled. (C) functional groups of fungi, (D) fungal orders, and (E) bacterial phyla affected significantly by plants growing in their own soil compared to growing in other soils in general manner, in plant functional group specific manner or plant species specific manner calculated using formula ln(home/away). The colors refer to plant species, green colors mark grasses and blue colors forbs. The effect of abundance in own soil vs other soils is calculated by dividing the taxa abundance in own soil with the abundance in all other soils. Lines mark when there is two times more or two times less members of fungi or bacteria in the soils.

Due to the importance of endophytes to plant performance we further investigated in the endophytic compartment, which microbial groups changed when the current plant was grown in conspecific soil, or in soils from a plant from the same family group. We detected that the relative abundance of potential fungal plant pathogens in the roots of grasses increased when grown in their own soil (log transformed relative abundance LME: F=13.84, p<0.001; Fig. 5C, Fig. S6). Forbs growing in grass soils exhibited higher relative abundances of plant pathogens and this was independent from whether the plants were grown in their own soil or in another soil (log transformed relative abundance LME: F=21.41, p<0.001, Fig. S6). Two fungal orders, Agaricales (LME: F=5.50, p=0.020) and Magnaporthales (LME: F=9.13, p=0.003), were enriched in roots of grass species that grew in their own soil, but this was not true for forbs growing in their own soil (i.e., a family specific soil effect; Fig. 5D). For the different grass species, different plant pathogens were enriched in roots when grown in their own soil (Fig. S7). The main pathogens that were enriched when grown in own soil compared to other soils were for *A. pratensis*, Magnaporthiopsis species, for *F. ovina, Alternaria* sp. and *Slopeiomyces cylindrosporus* and for *H. lanatus*, Magnaporthiopsis species and *Neoerysiphe nevoi*. For AMF we did not detect strong positive selection in roots of plants grown in their own soils and only some saprotrophic taxa such as Orbiliaceae, *Ceratobasidium* sp. and *Marasmius* sp. were affected by growing in their own soils (Fig. S8).

There were fewer *Actinobacteria* inside the roots of all plants when grown in conspecific compared to heterospecific soils (LME: F=4.01, p=0.047, Fig. 5E). Especially the abundance of *Streptomycetes* (own soil effect across species LME: F=5.95, p=0.016) and *Pseudonocardiaceae* (own soil effect across species LME: F=2.46, p=0.046; Fig. S10) were decreased when growing in own soil compared to growing in other soils across plant species. We detected family-specific own-soil effects for endophytic *Acidobacteria* (interaction own-soil*plant family: LME: F=7.55, p=0.007) and *Chloroflexi* (interaction own-soil*plant family LME: F=4.47, p=0.038) and species-specific responses to own and foreign soil for *Chlamydiae* (interaction own-soil*plant species LME: F=4.18, p<0.001) and *Elusimicrobia* (interaction own-soil*plant species LME: F=5.10, p<0.001). *Bradyrhizobium* (LME interaction own-soil*plant family: F=6.89, p<0.01) and *Acidibacter* (LME interaction own-soil*plant family: F=5.77, p=0.017) were increased if grasses were grown in their own soil, while *Pseudomonads* were four times more abundant in the roots of forbs (*J. vulgaris* or *T. officinale*) when grown in their own soil (Fig. S8). The relative abundance of the fungal class Helotiales was affected when grown in own soils, but in a plant species-specific manner (LME: F=2.95, p=0.014, Fig. 5D). For Helotiales, most sequences could not be assigned further than as *‘Helotiales* sp.’ and only three species that were present in more than 20 samples could be identified in more detail. The family responsible for the increase in relative abundance of Helotiales in *H. lanatus* soils was Hyaloscyphaceae that was only present in the roots if *H. lanatus* grew in its own soil. Furthermore, we detected that Pseudomonas and especially OTU83 was enriched in the roots of *J. vulgaris* and *T. officinale* when the plants were grown in their own soils (Dunn’s post hoc (home-away) JV: Z=2.38, p=0.024, TO: Z=2.27, p=0.046; fig S8).

## Discussion

We show that soil legacies from previous plants at species level and plant family level can be detected in the soil fungal community for at least five months after removal of the first plant and after subsequent colonization of the same soil by different plants. We find that the effect of the previous plant on soil fungal community structure can often be larger than the effect of the current plant species. This is important, as soils are dynamic and every soil arguably has a pre-existing legacy wrought by previous plants or plant communities. Our results indicate that when a plant arrives or is planted into the soil, even months after growing in this soil, it may still experience the microbial legacy effects created by the plants that grew previously in that soil. We can only speculate how long it will take before the legacy of the previous plant in the soil fungal community has disappeared entirely, as the five months of ‘current’ plant growth following one year of ‘previous’ plant conditioning was not enough for these soil legacy effects to fade away. This finding has important consequences for plant-growth experiments using field collected soils with previous legacies, but also for understanding plant community dynamics in natural and anthropogenic ecosystems, and suggests that the legacy of previous plants or plant communities on the soil microbiome lasts longer than thought previously.

For soil fungi, the effects of the previous plant on the soil community outweighed the effects of current plant, while for soil bacteria, each plant quickly modified its own microbiome although the influence of the previous plant was still detectable. These findings are in line with the conceptual idea that fungal growth rates are slower than those of bacteria (34,35) and that because of this, fungi are more stable and less affected by for instance temporal variability in the habitat or environment (31). Importantly, the more persistent effects of plants on the soil fungal communities than on bacterial communities may explain why most correlative studies that link plant responses and changes in the soil microbiome have shown that fungal communities drive plant community dynamics, while bacteria do not seem to strongly influence plant-soil-feedbacks (7,36) despite their known importance in rhizosphere processes (1,10). This is potentially due to faster turnover times of soil bacterial communities (34). Furthermore, due to their hyphal growth form, many fungi can simultaneously grow inside the roots and in the rhizosphere environment (37), and soils could therefore potentially predict intimate active fungal-plant relationships better than bacterial-plant relationships. Another possible option is that not all organisms detected with DNA based methods are active and thus part of the signal we are detecting originates from dead cells or inactive organisms (38). Alternatively, bacterial DNA could potentially be recycled quicker than fungal DNA which could be protected within the hyphae, but this needs further testing. However, recently it has been shown in the same ecosystem that approximately 80% of fungi detected in the rhizosphere with molecular DNA based methods also used here were actively participating in recycling plant-derived carbon (39), which further confirms the importance of soil fungi in soil-plant interactions.

Interestingly, the effects of the previous and current plant, especially on fungi, were conserved between the two plant families. Here, we show that even after five months, the fungal communities in current grass soils with a legacy of previous grasses grown on them differ from the fungal communities in the same grass soils with a legacy of previous forbs. Similarly, ‘current forb’ soils with a legacy of previous grasses on them had very different fungal communities than ‘current forb’ soils with a legacy of previous forbs. Other work has shown that plant family and functional group can explain a large portion of the variation in fungal community structure (31, 40, 41) and that this division may play a prominent role in plant community dynamics in natural grasslands (7,42). This is likely due to species in these groups having higher similarities within than between groups in terms of functional traits (43,44) and chemical composition (45, 46), which play an important role in shaping soil legacies. However, due to the higher phylogenetic variation observed in forbs (comprising many families) than in grasses (comprising one family), forbs generally exhibit a higher within-group variability than grasses in the modification of soil legacies. To balance our design for such phylogenetic patterns we used one plant family of either functional group in this study, this may limit the conclusions we can draw at the functional group level, although meta-analysis shows that the dichotomy generally has robust soil legacy effects (e.g.47).

One caveat of our study is that we did not measure soil chemistry mediated soil legacy effects shown to play a role mediating plant legacy effects in previous studies (13), although we have recently shown that the role of soil chemistry is minor under (similar) field conditions (7). Here, we related plant growth across plant species to soil fungal community structure and to the composition of endophytic microbiomes and especially plant pathogenic organisms. However, the soil abiotic parameters measured in this study, especially across plant species were rather low and we acknowledge that plant-soil feedback are driven by soil abiotic and biotic factors, often working also in synchrony (11,13). We could, however, pinpoint across plant species which organisms are likely drivers of the plant legacy effects and urge that further detailed investigations on their roles in plant-microbe-interactions are necessary.

An important outcome of our study is that soil legacies are taken up in early life stages and remain present inside the roots of the growing plants in a seemingly stable composition. This is supported by the observation that both the bacterial and the fungal endophyte community reflect the legacy of the previous plant. This may be important, as endophytic bacterial communities were more tightly linked to plant performance than soil bacterial communities, while for fungi both soil and endophytic communities influenced plant performance to a similar extent. This is probably due to the ability of fungi to grow as hyphae and thus bridging endophytic and soil environments (37). Previously, it has been shown that endophytic microbes are especially beneficial for plant growth through effects on plant nutrition, plant defense and interactions with the phytobiome, the response to abiotic constraints such as drought and on development (8–10,24). Here we show that the composition of the endophytic microbes is specific to the plant species carrying them. This could be partly explained by a strong influence of the inherited seed microbiome (25), which was reduced here by surface-sterilizing the seeds, but an effect of endophytic microbes of the seed cannot be ruled out. Our data indicate that the composition of the endophytic root compartment also depends on the soil legacy left by the previous plant that grew in the soil.

Root endophytic bacteria inhabiting the current plant were more sensitive to the soil legacies of the previous plant than soil and rhizosphere bacteria. This shows that bacteria are picked up from the soil in early plant ontogeny when the plant established in the new soil. Bacterial endophytes, and especially *Actinobacteria* and *Patescibacteria*, were modulated by the legacy of the previous plant, and influenced current plant growth. The role of *Patescibacteria* in plant health is still unclear, but they have been recently found inside the tissues of different plants (28,48). However, *Actinobacteria*, and especially *Streptomycetes*, are often detected inside plant roots and can act both as pathogens or beneficials to the plant (49, 50). Here, we show that root biomass increased across plant species when the relative abundance of *Actinobacteria* inside the roots increased. This correlation may suggest a generally positive role of this group of microbes in influencing plant root growth. Interestingly, we also show that all plant species had fewer *Actinobacteria*, and especially *Streptomycetes* inside their roots when grown in soils with a legacy of their own species. Previous studies indicated that monocultures may accumulate higher levels of antagonistic *Streptomycetes* in their soils (51) while, in contrast, our results indicate that these are depleted inside the roots when monocultures are grown. Recent work has shown how *Streptomyces coelicolor* shows an ‘intraspecific division of labour’, meaning a differentiation of functions adopted by cohorts within the same species, by generating heterogeneity within colonies at the chromosomal level (52). We speculate that plant species-specific selection on soil microbes, and specifically *Streptomycetes*, could drive the accumulation of antagonistic ‘strains’ through chromosomal heterogeneity, and this may provide an interesting avenue for further investigation on the role of these endophytic bacteria on conspecific plant-soil feedbacks.

We observed a trade-off between the AMF colonization inside plant roots and the respective root biomass. Plants with relatively fewer AMF in their roots had a higher belowground biomass, probably due to increased necessity to scavenge for nutrients and especially phosphorus (53). AMF have often been shown to explain (positive) plant-soil feedback (13,54), especially between plants that form arbuscular mycorrhizae and plants with other microbial mediated nutrient-acquisition strategies (55). We detected neither soil legacy-specific differences in AMF in the soil nor inside the roots of the plants, indicating that AMF may play only a minor role in plant-soil feedbacks in this specific system. It should be noted that the species that were used in this study, despite all forming arbuscular mycorrhizae, generally cause negative conspecific plant-soil feedbacks (15), which may be one reason for the absence of links between AMF and plant-soil feedback. We did detect after one month of growth lower relative abundance of AMF in soil of *J. vulgaris* when this plant was grown in its own soil compared to other soils, but this effect was transient and not detected in the roots.

There were strong negative feedbacks for all plant species. However, we observed only in grasses that this was due to accumulation of potential plant pathogens in the soil and inside roots when they were grown in soils with a legacy of their own species. The importance of the soil fungal community as a whole, and that of specific plant pathogenic fungi, has recently been shown to modulate plant community dynamics in grasslands (7,36,56). Here we show that grasses increase the abundance of plant pathogenic fungi in the soil when grown in monocultures. More importantly, we show that different grass species accumulate specific fungal pathogens that in turn cause negative plant-soil feedback when these grasses are grown in conspecific soils. In all six soil legacies, the soil fungal communities differed when a plant was grown in its own soil from the communities formed when grown in a soil with a legacy of another plant, and when this effect was largest, depended on plant species. Some of the plants in this experiment showed a stronger relationship with soil microbial communities at the beginning of the experiment, while for others the growth was more related to the current microbiome in the soils, still with a detectable legacy of previous microbes. The variability in the magnitude of plant-soil feedbacks between plant life stages has been noted earlier (16,18) and here we offer a microbial background on the observed phenomenon.

On the basis of our results, we propose a rethinking of how soil microbiome-mediated legacy effects work. First, no soil with plants growing on it is naive and without a legacy. Therefore, in order to predict what are the main factors predicting plant and especially crop growth, we need to look into the history of the soil and importantly also evaluate the plant ‘holobiome’ (57). We further show that part of the microbial legacy effects is plant-species specific while another part of the effects is plant family-specific. Especially, grasses generally had negative effects on all other grasses consequently growing in the same soils while mainly plant species-specific effects on forbs were detected. This advances our understanding on soil legacy effects and can be used also in agriculture for example in designing rotations to prevent accumulation of fungal plant pathogens. In a wider perspective, our results show that soil and root microbiomes are important for plant growth and that plants can be used to directionally change the microbiomes and hence steer plant growth. This can be done in an agricultural setting by selecting suitable cover crop species and using crop rotations. In conclusion, our study shows that soil legacies wrought by previous plants can remain present in the soil for months, even when subsequent plants colonize (and condition) the soil. However, our findings also suggest that plants take up microbes and especially bacteria from these pre-existing soil legacies in their endophytic compartments at a very early seedling stage, and that these may play a more prominent role in driving plant performance than the microbiome present around the roots of the older plants. Our study highlights also microbial species that consistently drive negative conspecific plant-soil feedbacks across plant-species, and characterizing the role of these species in plant-soil feedbacks will provide an exciting venue for further research.

## Materials and Methods

### Experimental design

The set-up of the conditioning phase is described in detail elsewhere (31) and shown in Figure 1. In short, thirty containers (48 cm x 80 cm x 50 cm) were filled with soil that was sieved through a 32 mm sieve. The soil was sourced from a grassland near Lange Dreef, Driebergen, The Netherlands (52°02′N, 5°16′E) and is characterized as holtpodzol, sandy loam (84% sand, 11% silt, 2% clay, ~3% organic matter, 5.9 pH, 1,151.3 mg/kg total N, 2.7 mg/kg total P, 91.0 mg/kg total K; analyses by Eurofins Analytico Milieu B.V., Barneveld, The Netherlands, using in-house methods; 58). Monocultures of ~100 individuals of six plant species were grown in these soils for one year (from May 2017 to May 2018). Each species was planted in a separate container in 5 replicate blocks in a randomized block design. We used three grass species (*Holcus lanatus* (HL), *Festuca ovina* (FO), *Alopecurus pratensis* (AP)) and three forb species (*Hypochaeris radicata* (HR), *Jacobaea vulgaris* (JV), and *Taraxacum officinale* (TO)), all of which are very abundant and commonly occur in former agricultural grasslands in the Netherlands. The experiment was conducted in the common garden at the Netherlands Institute of Ecology (NIOO-KNAW, Wageningen, The Netherlands, 51° 59′ N, 5° 40′ E). Mesocosms were watered regularly during the summer months to avoid desiccation.

In May 2018 the plants (‘previous plant’) were removed from the mesocosms and the mesocosms were divided into six smaller sections as shown in Figure 1 and supplementary video. Roots were not removed from the system to include the legacy effect of decomposing belowground biomass, as would be the case under natural conditions of plant removal (e.g. strong grazing or wild boar disturbance). Three weeks later, four seedlings of a single species grown from sterilized seeds on sterile glass beads were planted in each section. This equals to 180 experimental units (6 sections within 6 monocultures arranged in 5 blocks). Soil samples were collected using a small soil corer (12 cm deep, 7 mm diameter) from 4 locations in each section, pooled, homogenized, and immediately stored at −20 °C until molecular analysis. Samples were collected just prior to planting with the previous plants still in it (30 samples; May 2018 (33)), one month after planting (July 2018), after 3 months (September 2018), and after 5 months of plant growth (November 2018).

### Sample preparation and sequencing

The experiment was harvested in November 2018 and total oven-dried aboveground and belowground biomass was determined for each mesocosm. Samples from roots were collected from randomly selected washed root fragments, surface sterilized using published protocols (22,28) and stored in −20 °C prior to molecular analysis.

DNA from soil was extracted from 0.75 g of soil using the PowerSoil DNA Isolation Kit (Qiagen, Hilden, Germany) and from 0.5g of homogenized roots using MP Biomedical FastDNA™ Spin Kit following the manufacturer’s protocol. Fungal and bacterial DNA was amplified using primer sets ITS4ngs and ITS3mix targeting ITS2 region of fungi (59) and primers 515FB and 806RB targeting V4 rRNA region of bacteria (60–62), purified using Agencourt AMPure XP magnetic beads (Beckman Coulter). Adapters and barcodes were added to samples using Nextera XT DNA library preparation kit sets A-C (Illumina, San Diego, CA, USA). Separate equimolarly pooled libraries were constructed for bacteria and fungi. Bacterial samples (n=720) were analysed in 5 MiSeq runs (4 for bacteria in soils and 1 for bacteria in roots) and fungi in 3 MiSeq runs.

Libraries were sequenced using MiSeq PE250 at McGill University and Genome Quebec Innovation Center. Extraction negatives and a mock community, containing 10 fungal species, was included in each library and used to compare data between sequencing runs, detect possible contaminants and to investigate the accuracy of the bioinformatics analysis.

### Bioinformatic and statistical analysis

Bacterial and fungal sequences were analyzed using the DADA2 (63) and PIPITS (64) pipelines, respectively. For fungi, taxonomy was assigned using the rdp classifier against the UNITE fungal ITS database (65). Finally, the OTU table was parsed against the FunGuild (v1.1) database to assign putative life strategies to taxonomically defined OTUs (66) and this was further curated using in-house databases (67). All singletons and all reads from other than bacterial or fungal origin (i.e. plant material, mitochondria, chloroplasts and protists) were removed from the datasets. To account for large differences in read numbers, all samples with less than 1000 reads or more than 60 000 reads were removed which resulted in removal of 6 samples for bacteria and 24 samples for fungi. Furthermore, all OTUs and ASVs present in less than 5 samples with relative abundance of <0.001% were removed from the dataset. Cumulative sum scaling (CSS) was used to normalize the data (68,69). PERMANOVA model was constructed to investigate the effects of current and previous plant and the family of the current and previous plant on soil microbial community structure using Bray-Curtis distance in ‘vegan’ (70). A full model was run with all the time points to detect the effect of time with block as a strata and treatments nested in time but to answer main questions here, we also constructed simpler models with all the time points measured separately. To estimate the effect of growing in their own soil, also a PERMANOVA model was used with interaction own soil x plant identity/plant family as the main investigated factor. To investigate the sensitivity of individual plant species to microbiomes, also plant species specific PERMANOVAs were run for all time points. For all models run, betadisper was used to check the homogeneity of dispersion. We use the R^2^ values from the models used to estimate the amount of variation explained by a variable in the model. To visualize the effects of previous and current plant on microbial community structure, NMDS ordination (without further transformations) was used. To simplify the message, we chose to use for figures either NMDS1 or NMDS2 but always run the full model for statistical significance. ENVFIT with 999 permutations and block as strata was used to fit plant (shoot and root) biomass data as vectors to the community structure data. Relative abundances of bacterial phyla and fungal classes and orders were calculated. Furthermore, relative contributions of fungal functional guilds were calculated. All groups of bacteria and fungi present in less than 10 samples were removed from the analysis.

We tested the effect of previous plant and current plant and the plant family on plant biomass and relative abundances of bacterial and fungal taxa using a linear mixed effect model (package NLME) with block as a random factor. We did the testing independently for all time points and for root endophytes for the relative abundances. For all analysis we used posthoc Tukey HSD tests to test which treatments differed from each other. We explored for a normal distribution of residuals using QQ-plots and a Shapiro-Wilk test and homogeneity of variances using a Levene’s test. For the relative abundances of bacterial and fungal taxa, arcsin square root or ln transformations were used. The transformation of the data is mentioned in the text and in the figures. All p-values derived from multiple calculations (such as in figure 3D) were corrected with Benjamini & Hochenberg which relies on calculating the expected proportion of false discoveries among rejected hypotheses to control for false discovery rate (FDR) (71). For some of the individual OTUs if normality was not achieved, non-parametric tests such as Dunns post hoc was used to investigate the effects of growing in the own soil compared to other soils.

To calculate the difference in microbial community structure in own soil compared to away soils a model with ‘away’ plant species or its functional group as random factor was built and PERMANOVA was used. For the relative abundances and plant growth, linear mixed effect models were used with ‘away’ plant species as random factor with transformations described above. To calculate ‘home-away’ soil effects, formula ln(home/away) was used.

The R code of the analysis will be shared upon request.

## Supporting information

Data file (plant biomass)

## Acknowledgments

**General**: We thank Grace Wangsa Putra, Eefje Sanders, Anna Kielak and Stijn Hofhuis for help in harvesting the experiment and/or molecular work. The sequencing was done in collaboration with McGill University and Génome Québec Innovation Centre.

## Author contributions

SEH, RH, MH, KS, JRdL and TMB conceived the initial idea of the experiment, SEH, RH, MH, KS, JRdL and RJ set-up and harvested the experiment. SEH, KS and RJ performed the molecular work and SEH analysed the data. SEH, RH and TMB wrote the initial draft of the manuscript and all authors commented on it.

## Data and materials availability

Sequence data is can be found at ENA under accession number PRJEB38409. Plant growth data is submitted as a supplementary table.

