## Supplementary figures for "Persistence of plant-mediated microbial soil legacy effects in soil and inside roots"

**Supplementary Materials for** Persistence of plant-mediated microbial soil legacy effects in soil and inside roots

Fig. S1. The fungal community structure per previous and current plant species in all soils and across time.
Fig. S2. The bacterial community structure per previous and current plant species in all soils and across time
Fig. S3. The fungal community structure per plant species in all soils
Fig. S4. Fungal community structure affected by growing in own soils and in away soils at different times
Fig. S5. Bacterial community structure affected by growing in own soils and in away soils at different times.
Fig. S6. Potential plant pathogens and AMF in own soils and in soils of plants from other species and in grass and forb soils in time
Fig. S7. Fungi affected by growing in own soil.
Fig. S8. Bacteria affected by growing in own soil.
Movie S1. How mesocosms were divided into smaller mesocosms

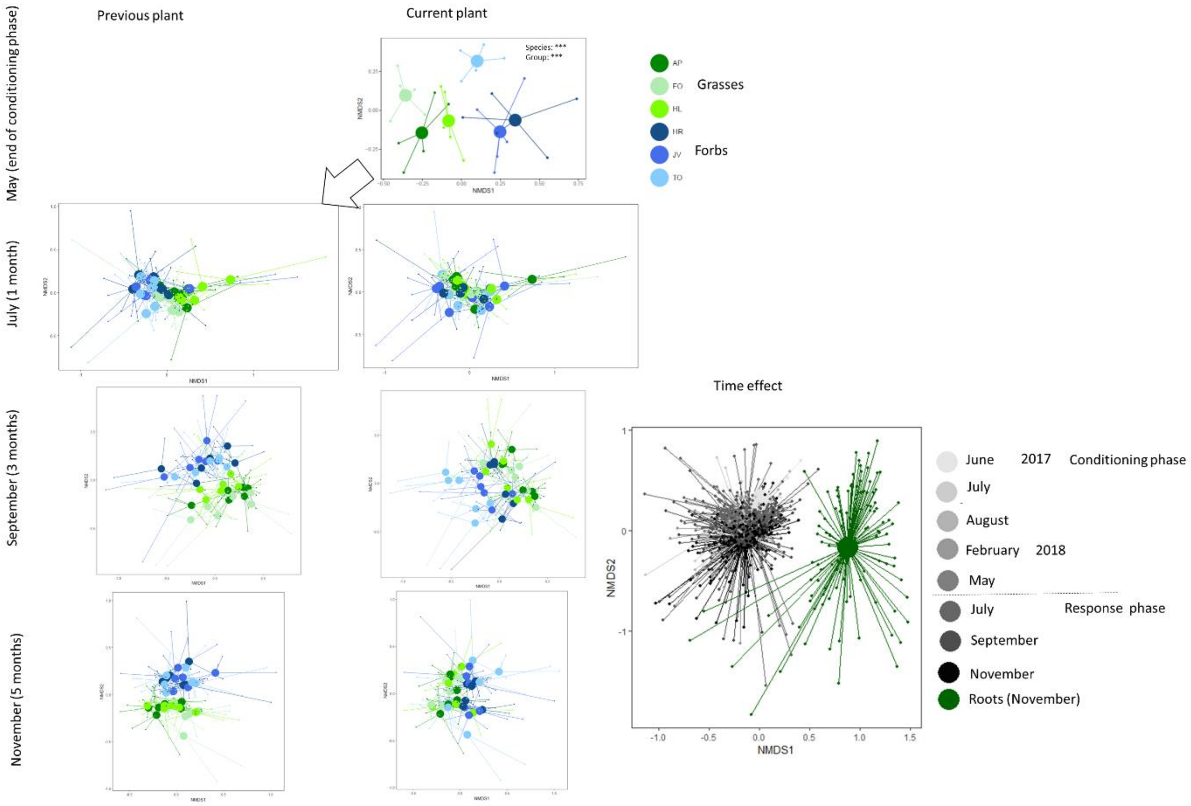


**Supplementary figure 1**. The fungal community structure per previous and current plant species in all soils and across time. The composition of fungal community at the end of the conditioning phase, and 1, 3 and 5 months after the start of the experiment depicted as centroids with variance (5 replicate containers) for each plant species with NMDS based on Bray Curtis dissimilarity. The mesocosms are colored based on conditioning (previous) plant species and by current plant species. The samples from the beginning are based on conditioning with current plant for 12 months and become the 'previous plant' for the new containers. The times are organized from light (June 2017; conditioning phase) to darker (November 2018; response phase) and contain data from the conditioning phase of the experiment (published earlier in Hannula et al. 2019) for comparison purposes. Root samples are marked in time series as green. The 2D stress values for end of conditioning phase is 0.18, for 1 month is 0.15, for 3 months 0.12 and for 5 months 0.14, and for the all data presenting variation in time 0.19.

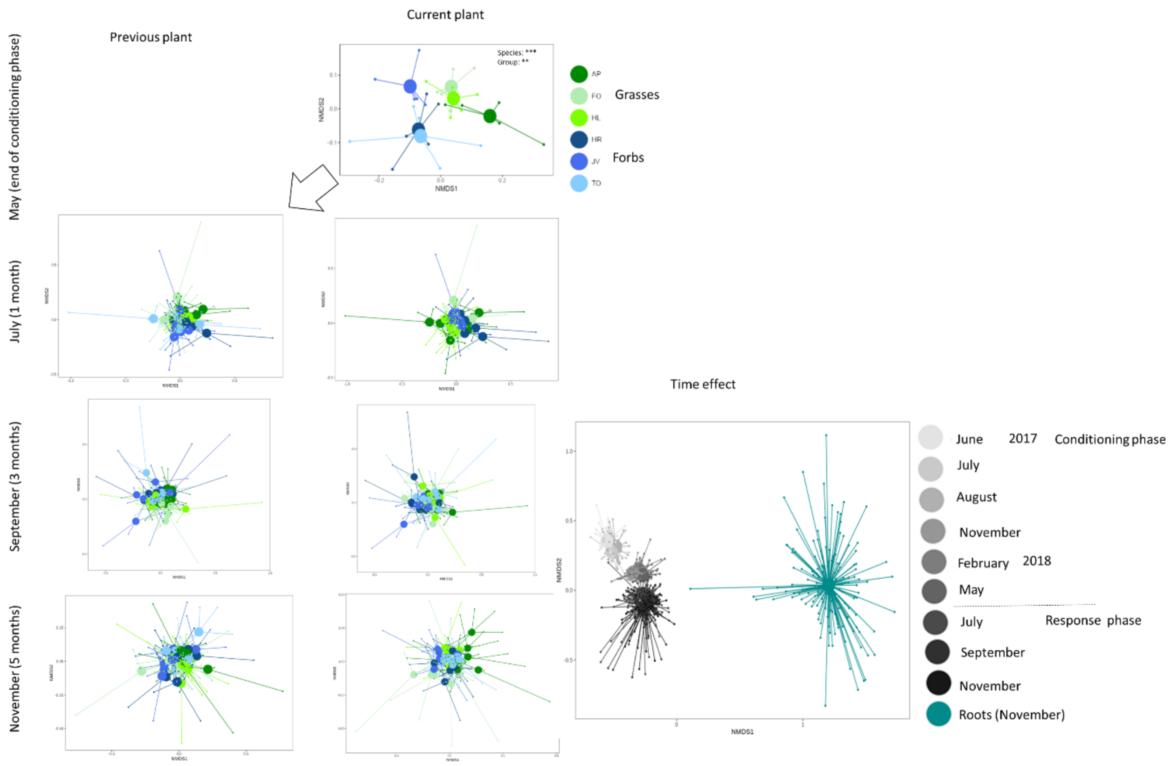


**Supplementary figure 2.** The bacterial community structure per previous and current plant species in all soils and across time. The composition of bacterial community at the end of the conditioning phase, and 1, 3, and 5 months after the start of the experiment depicted as centroids with variance (5 replicate containers) for each plant species with NMDS based on Bray Curtis dissimilarity. The mesocosms are colored based on conditioning (previous) plant species and by current plant species. The samples from the beginning are based on conditioning with current plant for 12 months and become the 'previous plant' for the new containers. The times are organized from light (June 2017; conditioning phase) to darker (November 2018; response phase) and contain data from the conditioning phase of the experiment (published earlier in Hannula et al. 2019) for comparison purposes. Root samples are marked with turqouise. The 2D stress values for end of conditioning phase is 0.14, for 1 month is 0.18, for 3 months 0.17 and for 5 months 0.15. The 2D stress value for the full time comparison was 0.21.

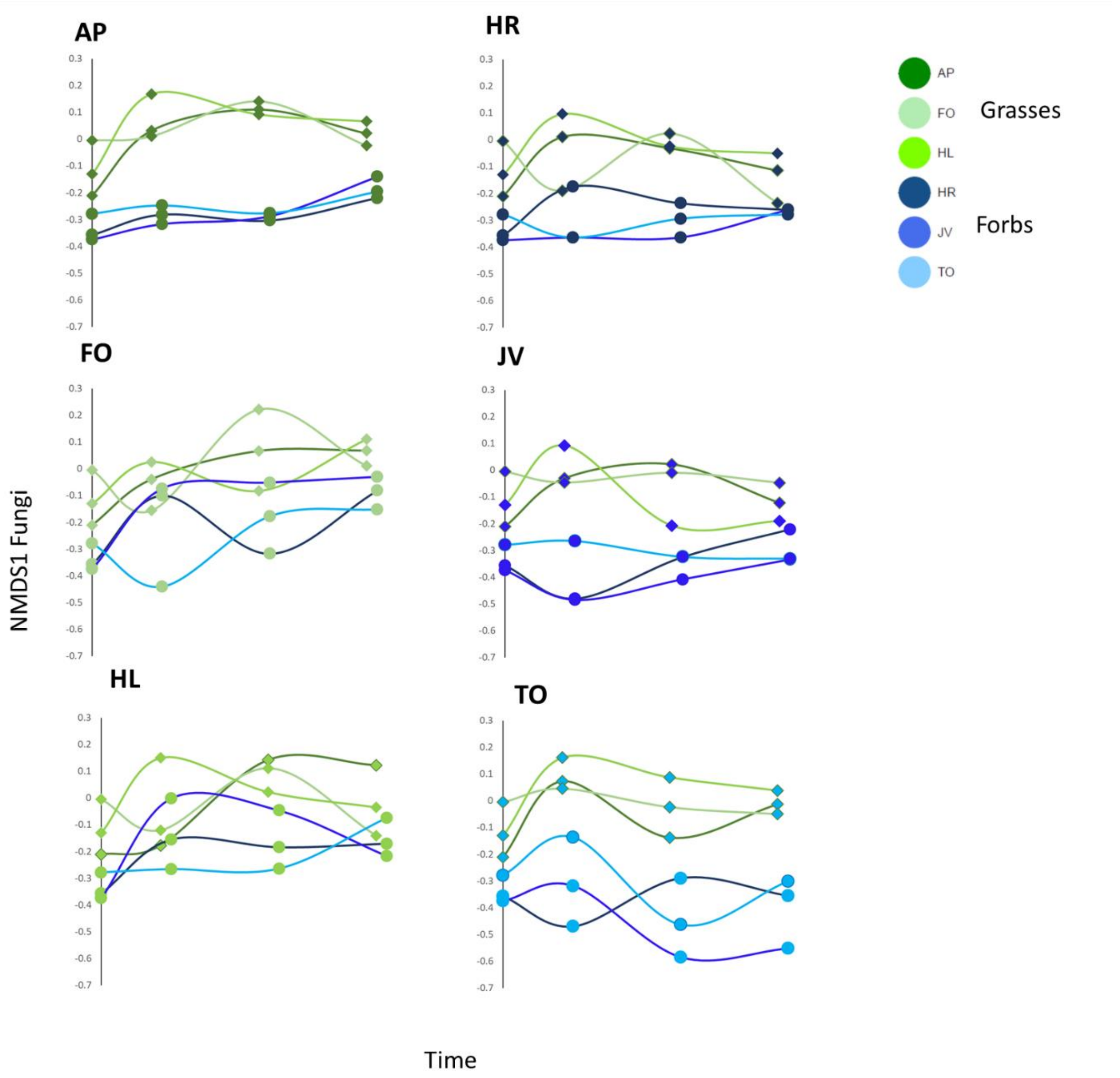


**Supplementary figure 3.** The fungal community structure per plant species in all soils. NMDS1 axis of fungal community per plant species (marker colors) and per soil types (line colors) as function of time. The NMDS is based on Bray Curtis dissimilarity and full data with variation is shown in Figure S2.

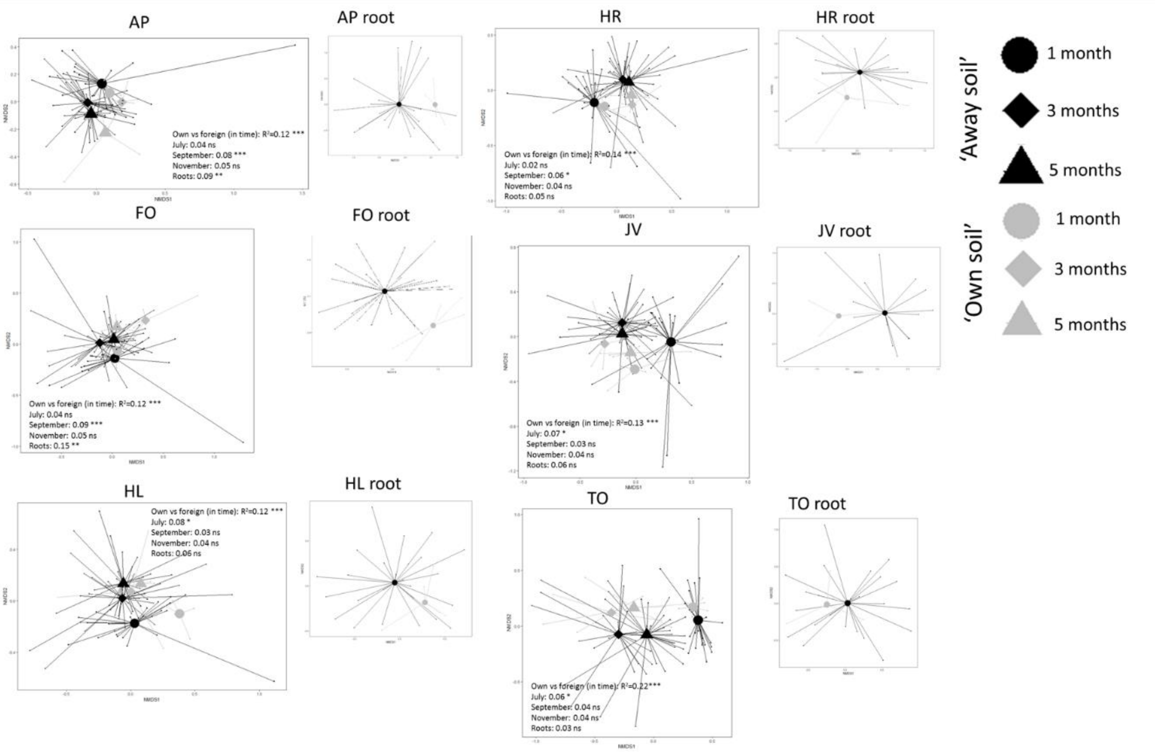


**Supplementary figure 4.** Fungal community structure affected by growing in own soils and in away soils at different times. Fungal communities in own (grey) and other soils from other plants (black) in time (symbols) per plant species. The fungal communities inside the roots are separately depicted. Statistical significance from PerMANOVA is given in the figure.

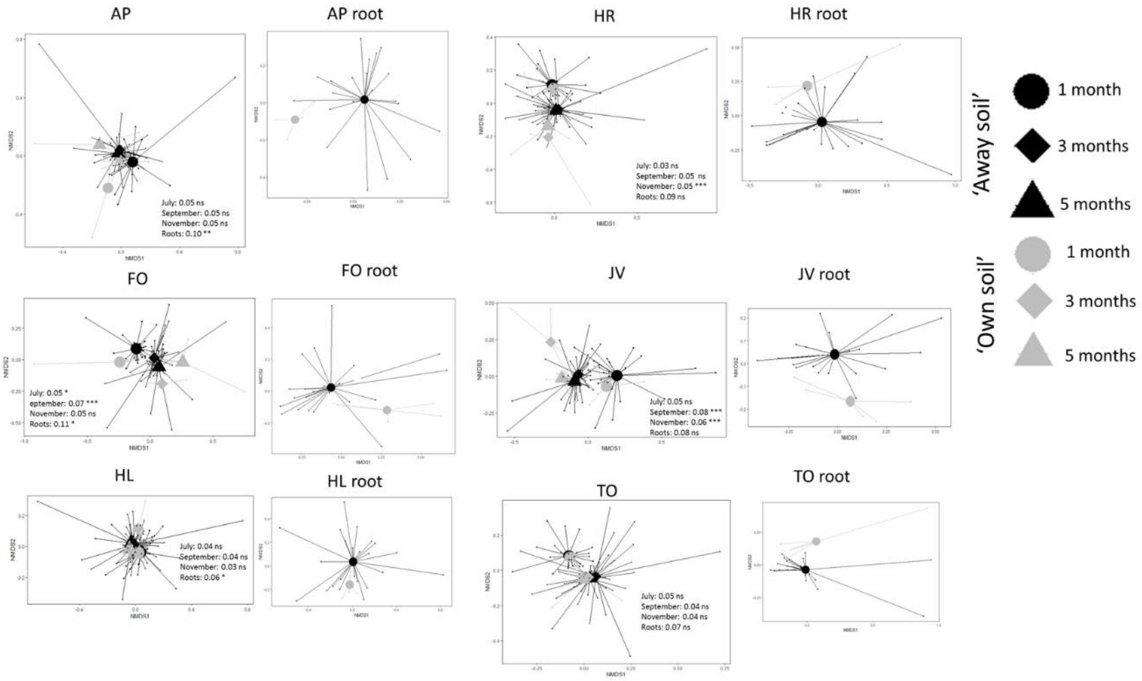


**Supplementary figure 5**. Bacterial community structure affected by growing in own soils and in away soils at different times. Bacterial communities in own (grey) and other soils from other plants (black) in time (symbols) per plant species. The bacterial communities inside the roots are separately depicted. Statistical significance from PerMANOVA is given in the figure.

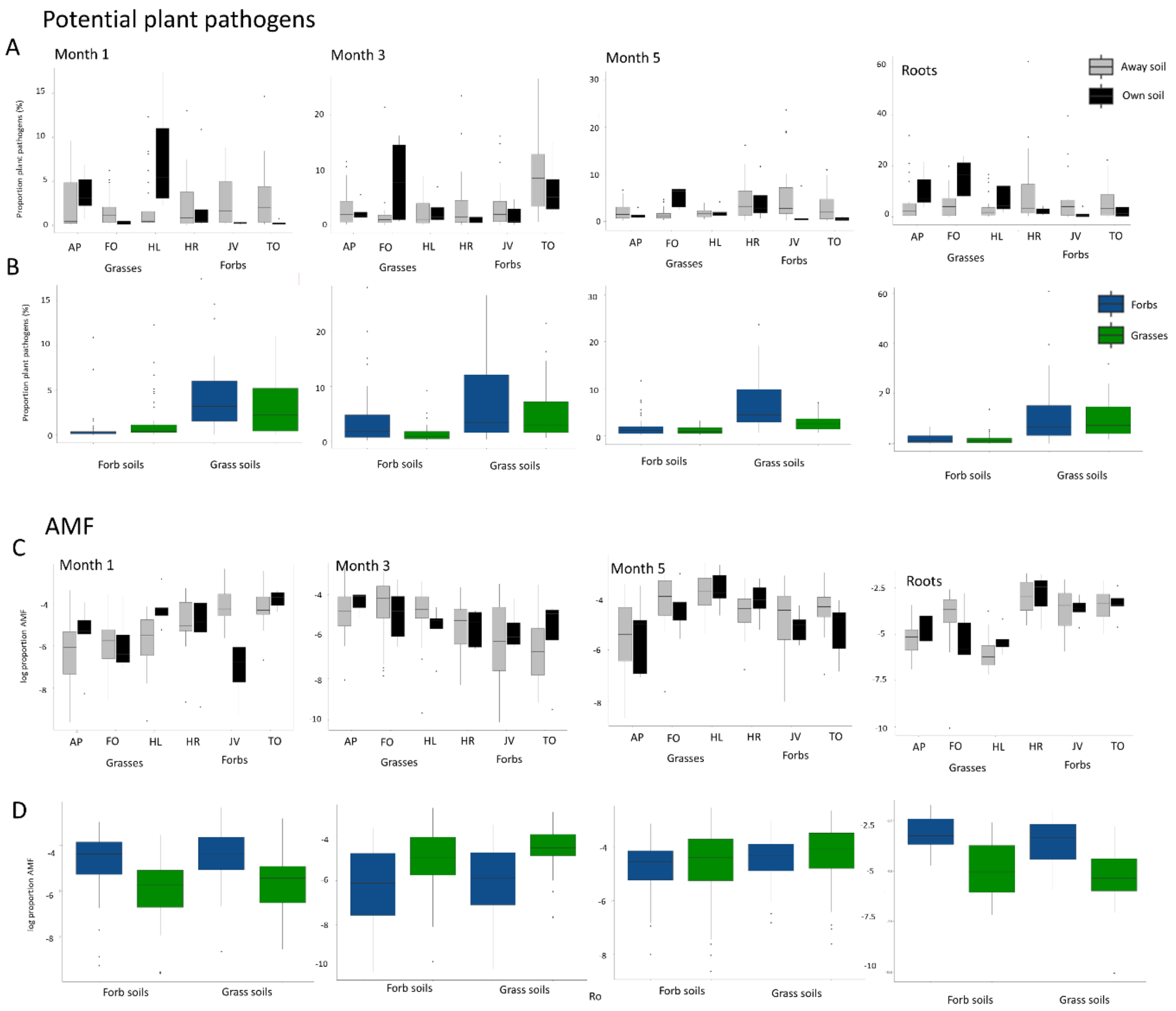


**Supplementary figure 6**. Potential plant pathogens and AMF in own soils and in soils of plants from other species and in grass and forb soils in time. Relative abundance of potential plant pathogens (A & B) and AMF (C & D) for all plants and time points when grown (A & C) in own soil (black) or away soils (grey) and plant functional groups grown in the soil of the same functional group or in the soil of another functional group. Blue colors denote forbs and green color grasses.

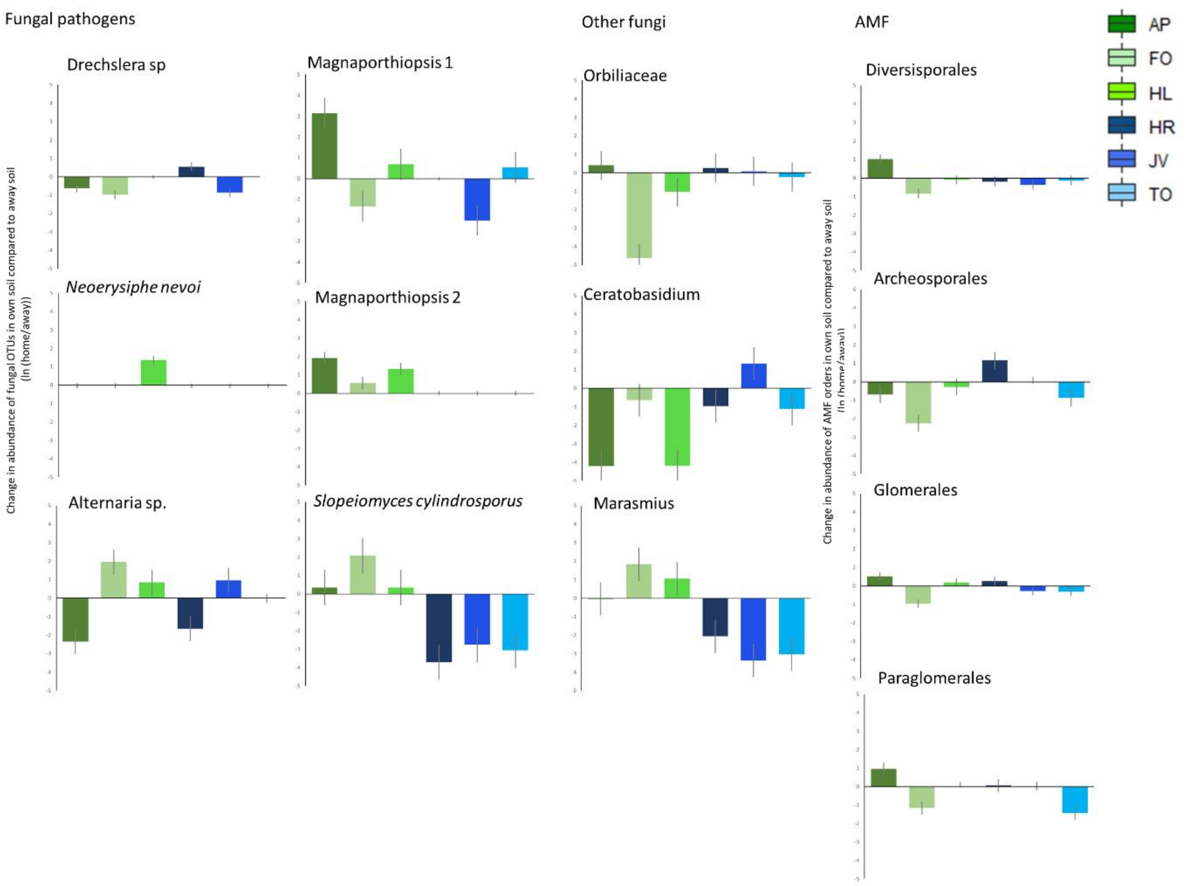


**Supplementary figure 7.** Fungi affected by growing in own soil. Phylotypes of fungal plant pathogens, and other fungi and orders of AMF inside the roots affected significantly by plants growing in their own soil compared to growing in another soil calculated using formula ln(home/away). Blue colors denote forbs and green colors grasses.

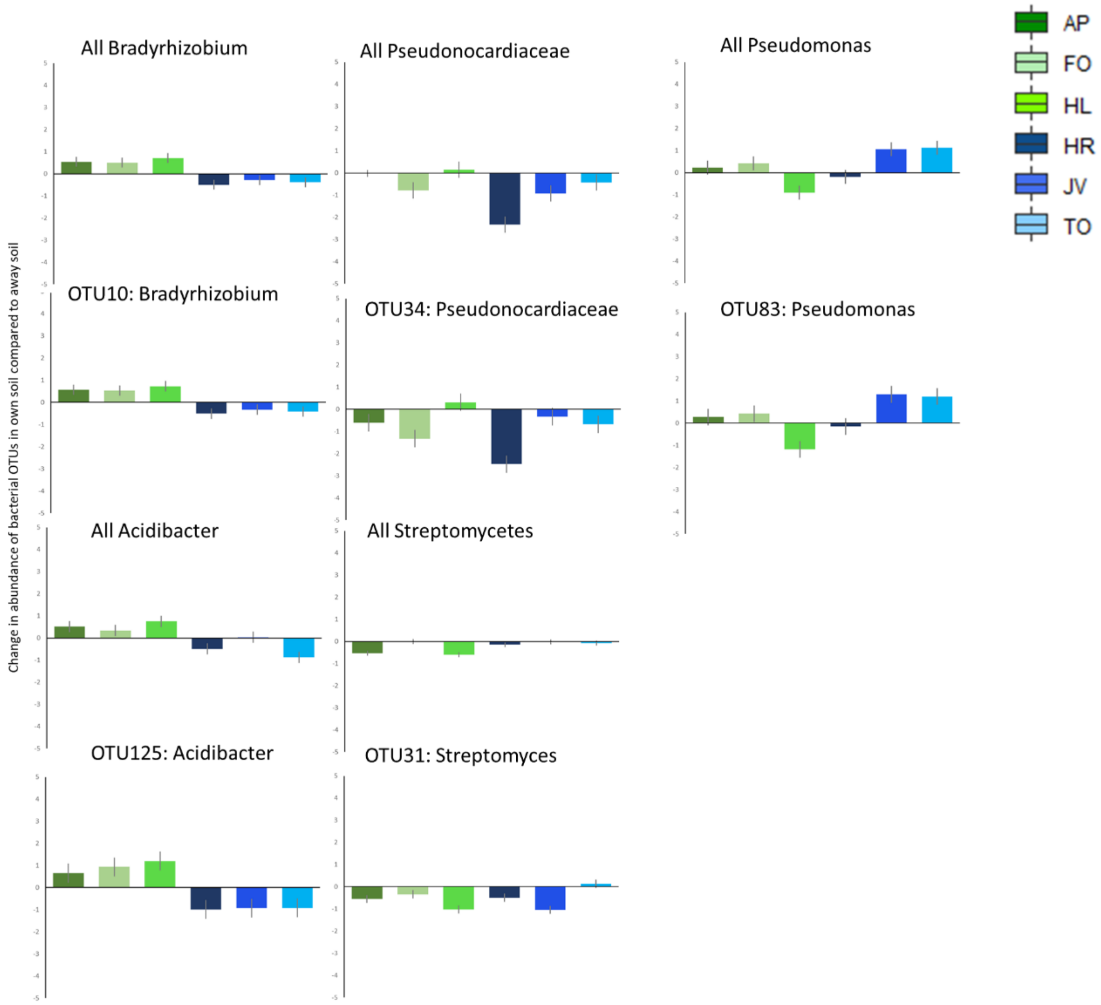


**Supplementary figure 8.** Bacteria affected by growing in own soil. Bacterial species and genera inside the roots affected significantly by plants growing in their own soil compared to growing in another soil calculated using formula ln(home/away). Blue colors denote forbs and green colors grasses.
